# The revised three-step detour pathway in dolichol biosynthesis is evolutionarily conserved in budding yeast

**DOI:** 10.64898/2026.04.13.718314

**Authors:** Kazuki Hanaoka, Kuya Matsunaga, Souichirou Shimizu, Soshi Sakai, Harald Pichler, Kouichi Funato

## Abstract

The identification of SRD5A3, a causative gene for congenital disorders of glycosylation (CDGs), together with its yeast ortholog *DFG10*, established the prevailing model that dolichol is synthesized from polyprenol in a single step. Subsequently, a recent discovery of DHRSX in CDG patients revised this view and led to the proposal of a three-step detour pathway for dolichol biosynthesis. However, it remains unclear whether this pathway represents a conserved mechanism or reflects evolutionary diversity in eukaryotes. Here, we identified *TDA5* as a yeast ortholog of DHRSX. Deletion of *TDA5* caused glycosylation defects, reduced dolichol levels, and accumulated polyprenol. All these phenotypes were rescued by expression of DHRSX, but not by *DFG10* or SRD5A3. These findings show that Tda5 serves the same function as DHRSX in yeast, thereby demonstrating conservation of the three-step detour pathway in yeast and supporting a broader eukaryotic framework for dolichol biosynthesis.

## Introduction

Dolichol is an essential long-chain polyisoprenoid lipid required for protein glycosylation in eukaryotic cells, and defects in its biosynthesis severely impair growth in budding yeast and cause congenital disorders of glycosylation (CDGs) in humans (1–5). Dolichol and sterols share farnesyl pyrophosphate as a common precursor in the mevalonate pathway, and defects in dolichol biosynthesis are known to affect sterol metabolism (2,6,7). As disruption of sterol metabolism is also linked to human developmental and neurological disorders (8,9), the consequences of impaired dolichol biosynthesis may extend beyond glycosylation defects. In the hitherto established model, dolichol is produced from polyprenol by a single reduction step, supported by identification of the human CDGs gene SRD5A3 and its yeast ortholog *DFG10* (3,5). However, dolichol remains detectable in cells lacking SRD5A3 or *DFG10*, suggesting that this model is incomplete and that additional enzymes remain to be identified (3). Recent work in human revised this pathway by proposing a three-step detour in which polyprenol is converted to dolichol through polyprenal and dolichal, with DHRSX catalyzing the first and third steps and SRD5A3 catalyzing the polyprenal-to-dolichal reduction step (10). Since no yeast ortholog of DHRSX has been identified, it remains unclear whether this revised pathway is conserved in yeast or widely conserved across eukaryotes.

## Result and Discussion

To identify a yeast ortholog of DHRSX, we focused on the short-chain dehydrogenase/reductase (SDR) superfamily, to which human DHRSX belongs (11). We screened deletion mutants of all 13 nonessential SDR genes in *Saccharomyces cerevisiae* for sensitivity to tunicamycin, an inhibitor of N-glycosylation (12). Among them, *tda5*Δ and *env9*Δ were hypersensitive to tunicamycin, although the primary functions of Tda5 and Env9 remain unclear (13). We therefore examined maturation of the vacuolar glycoprotein carboxypeptidase Y (CPY), a reporter for protein glycosylation during secretory pathway. Consistent with previous studies, *dfg10*Δ accumulated immature CPY (3), and *tda5*Δ also showed impaired CPY maturation (Fig.1*B*). We next asked whether these mutants exhibit sterol-related phenotypes associated with dolichol pathway defects, as impaired dolichol synthesis leads to squalene accumulation (2). As expected, *dfg10*Δ accumulated squalene. *tda5*Δ also showed a marked increase in squalene, whereas *env9*Δ showed only a modest increase (Fig.1*C*). Together with the CPY glycosylation defects, these findings support the idea that *TDA5* is more directly associated with dolichol synthesis than *ENV9*.

**Figure 1.**
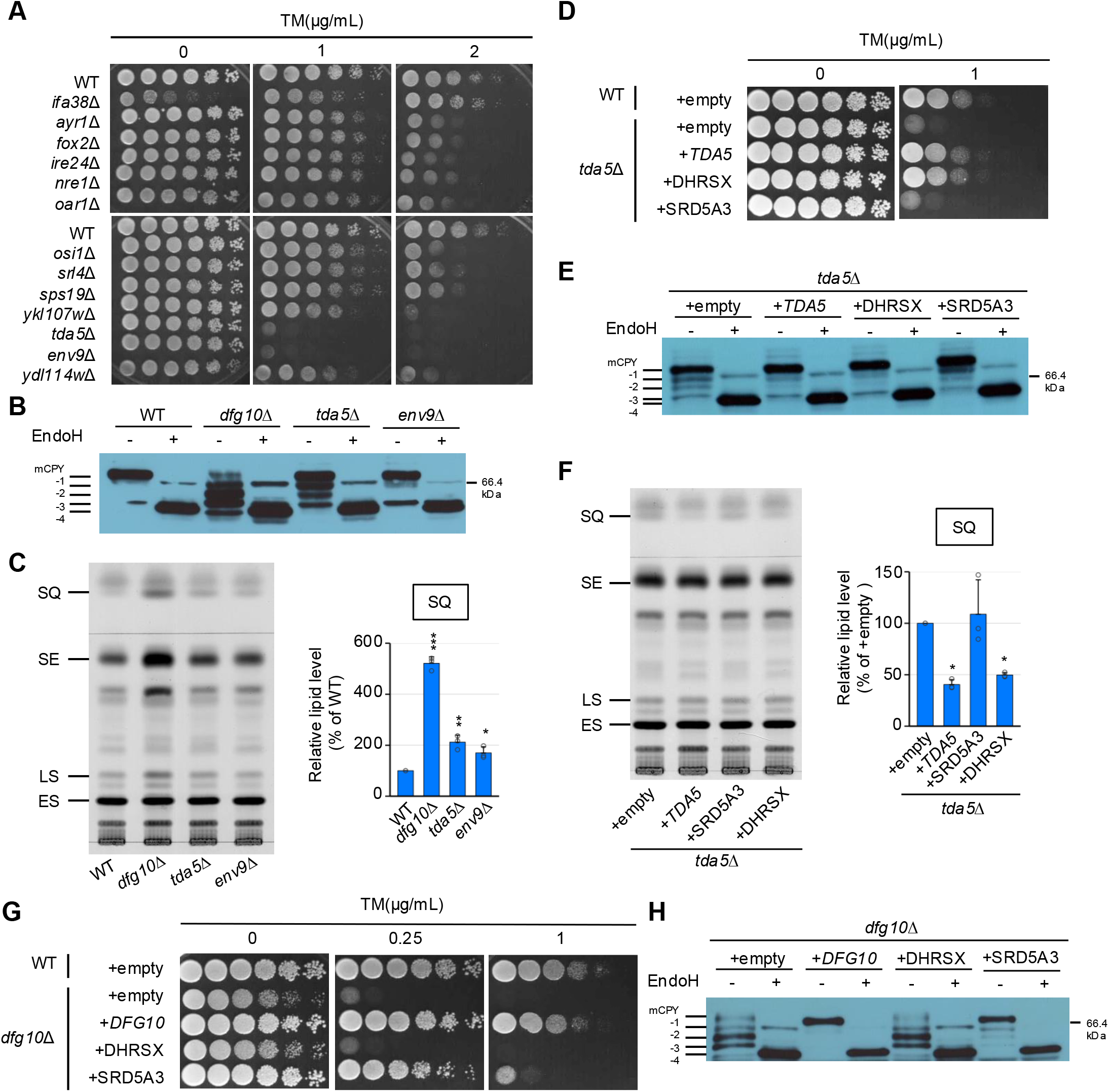
Screening of the SDR gene family identified*TDA5*, a functional yeast ortholog of human DHRSX. (A) Tunicamycin (TM) sensitivity assays of the indicated strains. (B) CPY glycosylation analyzed by immunoblotting with or without Endo H treatment. Mature CPY is indicated as mCPY, and hypoglycosylated immature CPY species are designated −1 to −4 from top to bottom. (C) TLC analysis of lipids from the indicated strains. SQ, squalene; SE, steryl ester; LS, lanosterol; ES, ergosterol. Graphs show SQ, SE, and LS levels relative to wild-type cells. (D, G) TM sensitivity assays of cells transformed with the indicated plasmids. (E, H) CPY glycosylation in cells transformed with the indicated plasmids. (F) TLC analysis of lipids from cells transformed with the indicated plasmids. Graph shows SQ levels relative to wild-type cells. *P < 0.05; **P < 0.01; ***P < 0.001.

We asked whether *TDA5* could be a functional ortholog of human DHRSX. Expression of human DHRSX, but not SRD5A3, from a CEN plasmid under the GPD promoter suppressed the tunicamycin sensitivity, abnormal CPY glycosylation, and squalene accumulation of *tda5*Δ cells (Fig.1*D-F*), suggesting that Tda5 serves the same function as DHRSX in yeast. Next, the fact that SRD5A3 expression, which can compensate for the function of Dfg10, the yeast ortholog of SRD5A3 (Fig.1 *G,H*), did not rescue the phenotypes of *tda5*Δ cells led us to confirm that Tda5 functions independently of Dfg10. As expected, the expression of *DFG10* and *TDA5* did not rescue the phenotypes of *tda5*Δ and *dfg10*Δ cells, respectively (Fig2.*A-D*). To further examine the relationship between *TDA5* and *DFG10*, we constructed a *tda5*Δ*dfg10*Δ double mutant. The double mutant showed more severe tunicamycin sensitivity than either single mutant, and its CPY glycosylation pattern combined features of both (Fig2.*E, F*). Together, these results suggest that *TDA5* and *DFG10* do not function solely in a simple linear order but may also act in parallel.

**Figure 2.**
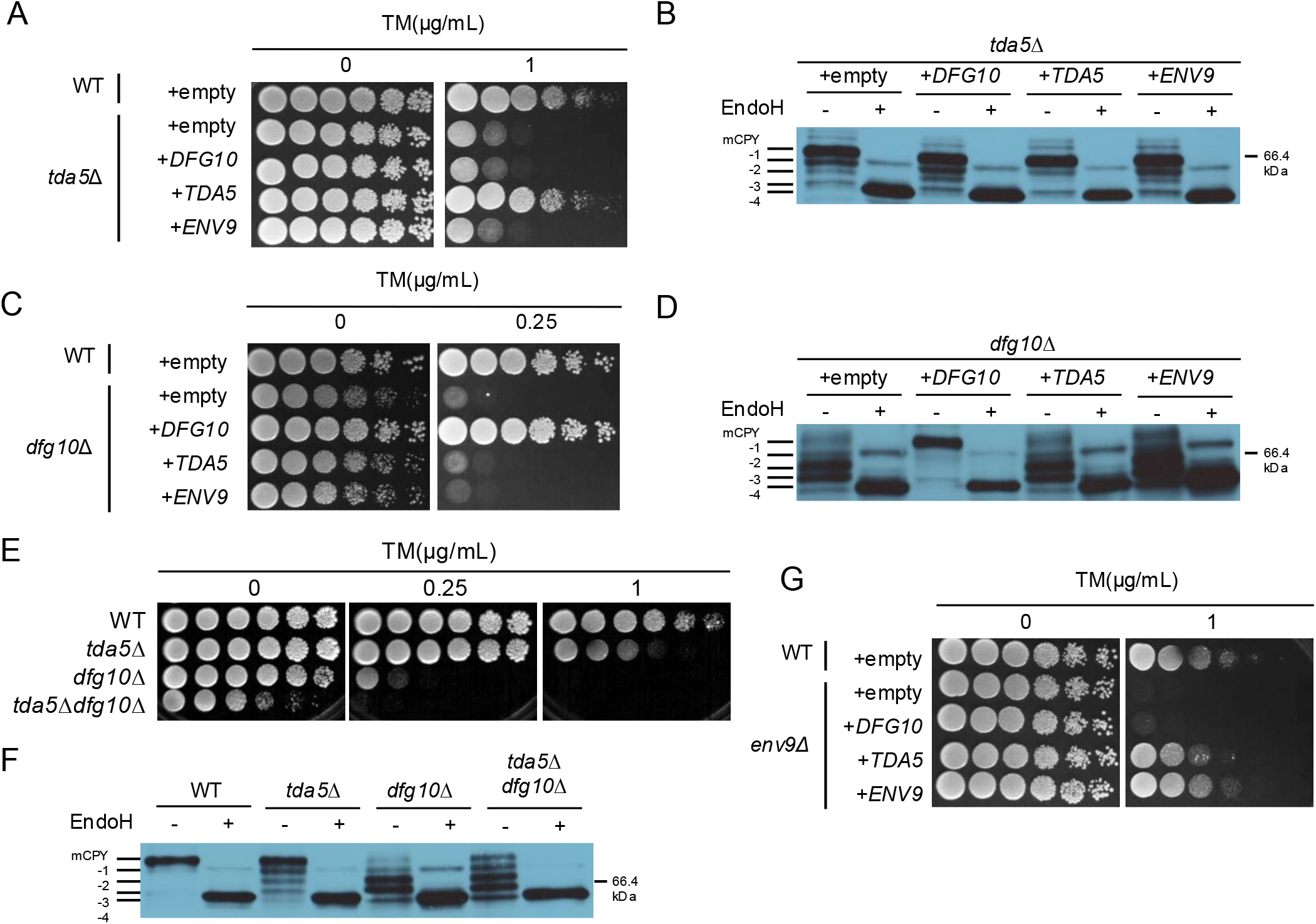
*TDA5* functions independently of *DFG10*, the yeast ortholog of human SRD5A3. (A, C, E, G) Tunicamycin (TM) sensitivity assays of cells transformed with or without the indicated plasmids, analyzed as in Fig. 1. (B, D, F) CPY glycosylation of cells transformed with or without the indicated plasmids, analyzed as in Fig. 1.

To directly evaluate the role of yeast orthologues in dolichol synthesis, we analyzed polyprenol and dolichol levels by TLC (2). In wild-type cells, dolichols were the predominant species and polyprenols were undetectable (Fig.3*A*). Consistent with previous reports (10), *dfg10*Δ cells showed increased polyprenol and decreased dolichol levels. In *tda5*Δ cells, polyprenol levels were significantly increased and dolichol levels were more severely reduced. By contrast, *env9*Δ cells showed only a slight reduction in dolichol levels, and not accumulation of polyprenols. Moreover, expression of *TDA5* or DHRSX restored both increased polyprenol and reduced dolichol levels in *tda5*Δ cells, whereas SRD5A3 did not (Fig.3*B*), further supporting *TDA5* as a functional ortholog of DHRSX. Remarkably, although polyprenol levels in the *tda5*Δ*dfg10*Δ double mutant were about twice those in *tda5*Δ cells, dolichol levels in the double mutant also increased by approximately two-fold (Fig.3*C*). This indicates that the polyprenol:dolichol ratio is the same between the two strains, arguing against simple recovery of dolichol synthesis, and instead, loss of *DFG10* may increase precursor flux into the dolichol pathway, thereby elevating both polyprenol and dolichol levels. This interpretation is consistent with the excessive accumulation of squalene in *dfg10*Δ cells (Fig.1*C*), and suggests that Dfg10 may be involved in flux distribution upstream of the dolichol and sterol branch (Fig.3*D*).

**Figure 3.**
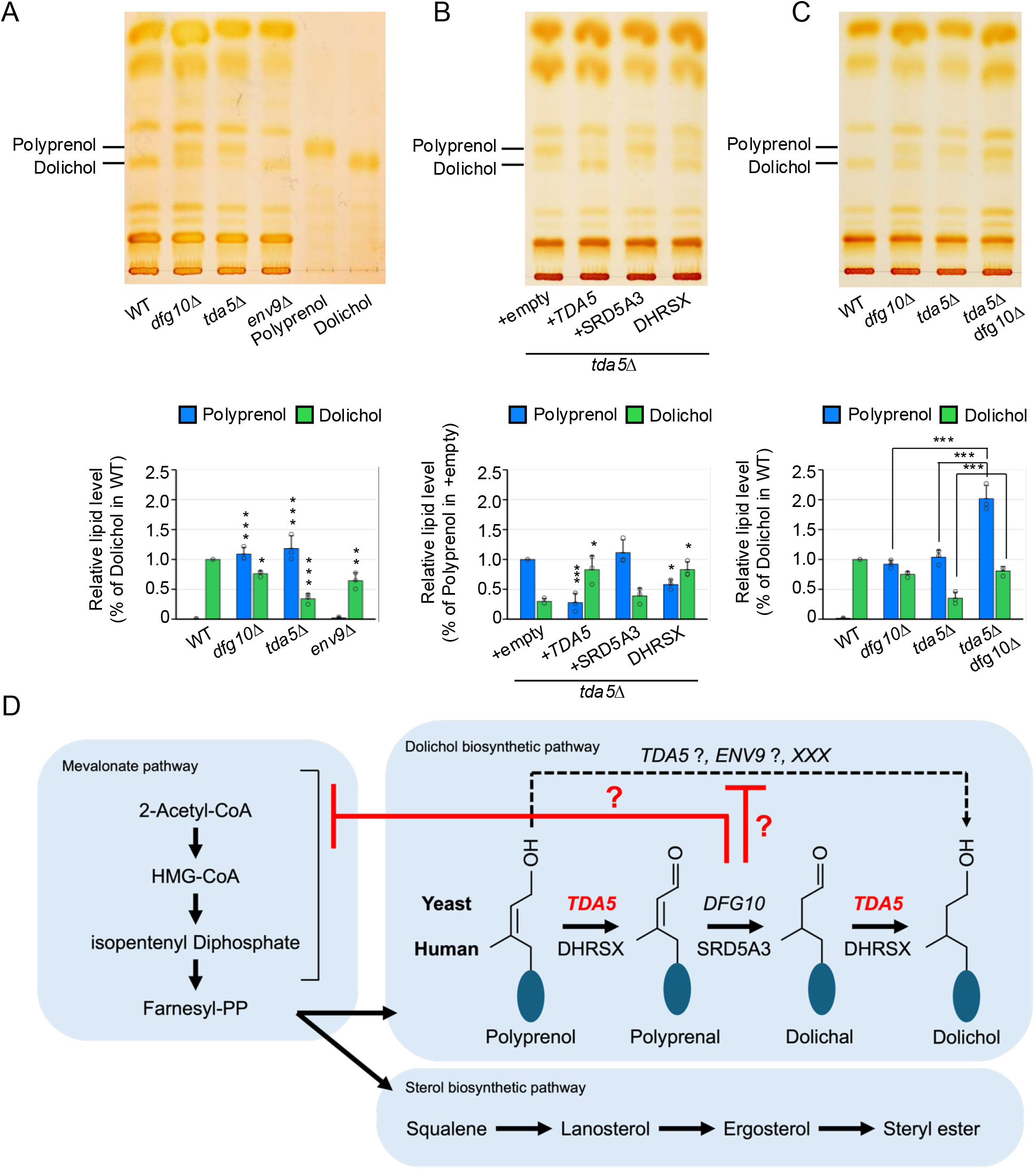
Loss of *TDA5* leads to increased polyprenol and decreased dolichol levels. (A–C) TLC analysis of polyprenol and dolichol in the indicated strains (A, C) or plasmid-transformed cells (B). Polyprenol and dolichol were assigned using authentic standards. (A). Graphs show dolichol normalized to wild type (A, C) or polyprenol normalized to the empty-vector control (B). Asterisks indicate significant differences for the indicated comparisons. ns, not significant; *P < 0.05, **P < 0.01, ***P < 0.001. (D) Proposed model for dolichol biosynthesis in budding yeast.

At the same time, persistence of dolichol in *dfg10*Δ cells indicates that dolichol can still be produced when the canonical three-step detour Tda5–Dfg10 pathway is disrupted, suggesting the existence of a bypass pathway. The bypass pathway may require Tda5 (Fig.3*D*), since Tda5 and Dfg10 appear to act in parallel in addition to linear order. Otherwise, Dfg10 may normally suppress the bypass pathway, and if Dfg10 function is impaired, the pathway may become activated. As dolichol remained detectable even in the *tda5*Δ*dfg10*Δdouble mutant, this bypass cannot be explained by the action of Tda5 alone and other factors are likely involved (Fig.3*D*). One candidate is Env9, because expression of *TDA5* suppressed the tunicamycin sensitivity of *env9*Δ(Fig.2*G*), whereas *ENV9* did not rescue the phenotypes in *tda5*Δ (Fig.2*A, B*), suggesting that Env9 may retain only a subset of Tda5 functions. Alternatively, a yet unidentified factor may contribute to this bypass.

Finally, the findings presented here reveal that Tda5 is a functional yeast ortholog of human DHRSX and support the evolutionary conservation of the revised three-step detour route from polyprenol to dolichol, and suggest that budding yeast may retain the bypass one-step pathway for direct conversion of polyprenol to dolichol.

## Materials and Methods

Details of the strains, plasmids, and experimental methods used in this study are provided in the Supporting Information.

## Acknowledgments

This work was supported by the Japan Society for the Promotion of Science (JSPS), Grants-in-Aid for Scientific Research (KAKENHI), Japan (21K19088) to K. F.

## Author Contribution

K.H., K.M., S.S.,and S.S.: Investigation, data curation and formal analysis.K.H., H.P. and K.F.: Conceptualization and writing

## References

1. M. Aebi, N-linked protein glycosylation in the ER. Biochim. Biophys. Acta 1833, 2430–2437 (2013).

2. M. Sato et al., The yeast RER2 gene, identified by endoplasmic reticulum protein localization mutations, encodes cis-prenyltransferase, a key enzyme in dolichol synthesis. Mol. Cell. Biol. 19, 471–483 (1999).

3. V. Cantagrel et al., SRD5A3 is required for converting polyprenol to dolichol and is mutated in a congenital glycosylation disorder. Cell 142, 203–217 (2010).

4. E. J. Park et al., Mutation of Nogo-B receptor, a subunit of cis-prenyltransferase, causes a congenital disorder of glycosylation. Cell Metab. 20, 448–457 (2014).

5. J. Denecke, C. Kranz, Hypoglycosylation due to dolichol metabolism defects. Biochim. Biophys. Acta 1792, 888–895 (2009).

6. E. Currie et al., High confidence proteomic analysis of yeast LDs identifies additional droplet proteins and reveals connections to dolichol synthesis and sterol acetylation. J. Lipid Res. 55, 1465–1477 (2014).

7. K. Grabińska, G. Palamarczyk, Dolichol biosynthesis in the yeast Saccharomyces cerevisiae: An insight into the regulatory role of farnesyl diphosphate synthase. FEMS Yeast Res. 2, 259–265 (2002).

8. F. D. Porter, G. E. Herman, Malformation syndromes caused by disorders of cholesterol synthesis. J. Lipid Res. 52, 6–34 (2011).

9. F. M. Platt et al., Disorders of cholesterol metabolism and their unanticipated convergent mechanisms of disease. Annu. Rev. Genomics Hum. Genet. 15, 173–194 (2014).

10. M. P. Wilson et al., A pseudoautosomal glycosylation disorder prompts the revision of dolichol biosynthesis. Cell 187, 3585–3601.e22 (2024).

11. G. Zhang et al., DHRSX, a novel non-classical secretory protein associated with starvation induced autophagy. Int. J. Med. Sci. 11, 962–970 (2014).

12. A. D. Elbein, Glycosylation inhibitors for N-linked glycoproteins. Methods Enzymol. 138, 661–709 (1987).

13. I. M. Siddiqah et al., Yeast ENV9 encodes a conserved lipid droplet (LD) short-chain dehydrogenase involved in LD morphology. Curr. Genet. 63, 1053–1072 (2017).

14. H. Sagami et al., Enzymatic formation of dehydrodolichal and dolichal, new products related to yeast dolichol biosynthesis. J. Biol. Chem. 271, 9560–9566 (1996).

