## Supporting information for "The revised three-step detour pathway in dolichol biosynthesis is evolutionarily conserved in budding yeast"

**This PDF file includes:**

Extended Materials and Methods

Supporting Information References

**Extended Materials and Methods**

**Lists of the strains**

| Strain No. | Description | Genotype | Source | Corresponding Figure No. |
| --- | --- | --- | --- | --- |
| FKY7486 | WT | Mat a, ura3 his3 leu2 met15 lys2 | This study | Fig. 1, 2, 3 |
| FKY8188 | ifa38Δ | ura3 his3 leu2 ifa38Δ;;KanMX | This study | Fig. 1 |
| FKY7195 | ayr1Δ | Mat a, ura3 his3 leu2 lys2 MET15 ayr1Δ;;KanMX | This study | Fig. 1 |
| FKY7236 | fox2Δ | Mat a, ura3 his3 leu2 LYS2 met15 fox2Δ;;KanMX | This study | Fig. 1 |
| FKY7256 | irc24Δ | Mat a, ura3 his3 leu2 LYS2 met15 irc24Δ;;KanMX | This study | Fig. 1 |
| FKY7274 | nre1Δ | Mat a, ura3 his3 leu2 lys2 met15 nre1Δ;;KanMX | This study | Fig. 1 |
| FKY7289 | oar1Δ | Mat a, ura3 his3 leu2 LYS2 MET15 oar1Δ;;KanMX | This study | Fig. 1 |
| FKY7304 | osi1Δ | Mat alpha, ura3 his3 leu2 lys2 met15 osi1Δ;;KanMX | This study | Fig. 1 |
| FKY7320 | srl4Δ | Mat a, ura3 his3 leu2 lys2 met15 srl4Δ;;KanMX | This study | Fig. 1 |
| FKY7339 | sps19Δ | Mat alpha, ura3 his3 leu2 lys2 met15 sps19Δ;;KanMX | This study | Fig. 1 |
| FKY7356 | ykl107wΔ | Mat a, ura3 his3 leu2 LYS2 MET15 ykl107wΔ;;KanMX | This study | Fig. 1 |
| FKY7513 | tda5Δ | Mat a, ura3 his3 leu2 MET15 lys2 tda5Δ;;KanMX | This study | Fig. 1, 2, 3 |
| FKY7630 | env9Δ | Mat alpha, ura3 his3 leu2 MET15 lys2 env9Δ;;KanMX | This study | Fig. 1, 2, 3 |
| FKY7638 | ydl114wΔ | ura3 his3 leu2 MET15 lys2 ydl114wΔ;;KanMX | This study | Fig. 1 |
| FKY7988 | dfg10Δ | Mat a, ura3 his3 leu2 MET15 dfg10Δ;;KanMX | This study | Fig. 1, 2, 3 |
| FKY8091 | WT | ura3 his3 leu2 | This study | Fig. 2 |
| FKY8092 | tda5Δ | ura3 his3 leu2 tda5Δ;;KanMX | This study | Fig. 2 |
| FKY8093 | dfg10Δ | ura3 his3 leu2 dfg10Δ;;KanMX | This study | Fig. 2 |
| FKY8094 | tda5Δdfg10Δ | ura3 his3 leu2 tda5Δ;;KanMX dfg10Δ;;KanMX | This study | Fig. 2 |

**Lists of the plasmids**

| Plasmid No. | Name | Marker | Promoter | Source | Description | Corresponding Figure |
| --- | --- | --- | --- | --- | --- | --- |
| FKP  22 | pRS416-GPD | URA3 | GPD | Mumberg et al., Gene, 1995 | empty | Fig. 1, 2, 3 |
| FKP  1234 | pRS416-GPD *TDA5* | URA3 | GPD | This study | The *TDA5* coding region was PCR-amplified from yeast genomic DNA and ligated into the BamHI-XhoI sites. | Fig. 1, 2, 3 |
| FKP  1278 | pRS416-GPD *DFG10* | URA3 | GPD | This study | The *DFG10* coding region was PCR-amplified from yeast genomic DNA and ligated into the BamHI-XhoI sites. | Fig. 1, 2, 3 |
| FKP  1303 | pRS416-GPD DHRSX | URA3 | GPD | This study | The *DHRSX* coding region was PCR-amplified from human cDNA library and ligated into the BamHI-XhoI sites. | Fig. 1, 2, 3 |
| FKP  1315 | pRS416-GPD SRD5A3 | URA3 | GPD | This study | The *SRD5A3* coding region was PCR-amplified from human cDNA library and ligated into the XbaI-XhoI sites. | Fig. 1, 2, 3 |
| FKP  1366 | pRS416-GPD *ENV9* | URA3 | GPD | This study | The *ENV9* coding region was PCR-amplified from yeast genomic DNA and ligated into the BamHI-XhoI sites. | Fig. 1, 2, 3 |

**Serial dilution assays for tunicamycin sensitivity**

Yeast cells, without or with plasmids, were cultured in rich YP medium (1% yeast extract, 2% peptone) supplemented with 0.2% adenine and containing 2% glucose (YPD) as carbon source at 25℃, and adjusted to OD_600_ 1.0 in sterile water. Fivefold serial dilutions were spotted onto fresh YPD plates containing tunicamycin at the concentrations indicated in the figure legends and incubated at 25℃ for 3 days.

**Immunoblot analysis of CPY glycosylation**

Yeast cells were grown in YPD at 25℃ to early log phase and lysed with glass beads in SDS-containing buffer. After removal of cell debris, lysates were treated with or without Endo-H at 37℃ for 60min, separated by SDS/PAGE, transferred to nitrocellulose membranes, and analyzed by immunoblotting with anti-CPY antibody. Signals were detected using HRP-conjugated secondary antibody and ECL substrate (1). Endo-H treatment was performed as described previously (2).

**TLC analysis of neutral lipids and polyprenol/dolichol**

Yeast cells were grown in YPD at 25℃ to early log phase, collected, washed, and adjusted to an OD_600_ of 10. Total lipids were extracted using chloroform-methanol-water (CMW, 10:10:3, v/v/v). For analysis of neutral lipids, extracts were separated by thin-layer chromatography (TLC) using petroleum ether-diethyl ether-acetic acid (25:25:1, v/v/v) for the first third of the plate, followed by petroleum ether-diethyl ether (49:1, v/v) for the remaining distance. Lipids were visualized by staining with MnCl2 reagent [0.63 g MnCl_2_·4H_2_O dissolved in methanol-water-sulfuric acid (60:60:4, v/v/v)] for 10 s (3,4). For analysis of polyprenol and dolichol, lipid extracts were separated by TLC using benzene-ethyl acetate (95:5, v/v) and visualized by exposure to iodine vapor for 15 min (5). Lipid bands were quantified using ImageJ.

**Statistical analysis**

All statistical analyses and graphing were performed using R (R Foundation for Statistical Computing, Vienna, Austria). Comparisons among multiple groups were performed by one-way ANOVA followed by Tukey’s multiple comparison test, as indicated in the figure legends. Data are presented as mean ± S.D. unless otherwise stated. In the graphs, bars indicate the mean, error bars indicate S.D., and dots indicate values from three independent biological replicates. Statistical significance is indicated as ns, not significant; *, p < 0.05; **, p < 0.01; ***, p < 0.001.

**Supporting Information References**

1. K. Kajiwara et al., Osh proteins regulate COPII-mediated vesicular transport of ceramide from the endoplasmic reticulum in budding yeast. J. Cell Sci. 127, 376–387 (2014).
2. N. Kageyama-Yahara, H. Riezman, Transmembrane topology of ceramide synthase in yeast. Biochem. J. 398, 585–593 (2006).
3. A. Ikeda et al., Tricalbins are required for non-vesicular ceramide transport at ER-Golgi contacts and modulate lipid droplet biogenesis. iScience 23, 101603 (2020).
4. Y. Yang et al., Retrograde Golgi-to-ER transport is regulated by diacylglycerol in Saccharomyces cerevisiae. J. Cell Sci. jcs264537 (2026).
5. M. Sato et al., The yeast *RER2* gene, identified by endoplasmic reticulum protein localization mutations, encodes cis-prenyltransferase, a key enzyme in dolichol synthesis. Mol. Cell. Biol. 19, 471–483 (1999).
